# A new method for the sampling and preservation of placental specimens in low-resource settings for the identification of *P. falciparum* and analysis of nucleic acids

**DOI:** 10.1101/2022.01.12.476059

**Authors:** Patrick S. Potoczak, Beverly I. Strassmann, Claudius Vincenz

## Abstract

Collection, preservation, and shipment of histological specimens in low-resource settings is challenging. We present a novel method that achieved excellent preservation of placental specimens from rural Mali by using formalin fixation, ethanol dehydration, and long-term storage in a solar-powered freezer. Sample preservation success was 92%, permitting evaluation of current and past malaria infection, anemia, placental maturity, and inflammation. Using RNAscope^®^ hybridization we were able to visualize cell-specific gene expression patterns in the formalin-fixed paraffin-embedded (FFPE) specimens. Additionally, our method entailed mirrored sampling from the two cut faces of a cotyledon, one for the FFPE workflows and the other for storage in RNAlater™ and RNA-seq.

## Introduction

Neutral-buffered formalin is routinely used for tissue preservation including for the placenta [1–5]. For this project, we sought to preserve placentas to detect malaria infection. Formaldehyde can produce artifacts that resemble hemozoin deposits, the waste product of the malaria parasite, particularly when acid formalin is used [6]. Even specimens fixed with neutral formalin can be subject to artifacts under conditions of prolonged fixation, elevated temperature, and high humidity [7]. We collected placental specimens in a rural location in Mali, West Africa, that is endemic for malaria (*Plasmodium falciparum*) and where ambient temperature often exceeds 40°C. We report a new collection protocol for placental specimens collected following delivery that minimized the formalin artifacts, required little on site infrastructure, and resulted in specimens that could be shipped on dry ice. Our collection and preservation protocol routinely produced high quality histological specimens suitable for the assessment of morphological features of the delivered placenta, including changes associated with active or past placental malaria infection, and suitable for *in situ* RNA hybridization. In parallel, we also collected specimens in RNAlater™ without fixation for molecular analysis of nucleic acids.

### 2.1. Collection and processing

The placentas came from women who participated in a longitudinal study on the Bandiagara Escarpment in Mali. The cohort and IRB approval is described in detail in Vincenz et al. [8, 9]. To obtain specimens from term placentas, we identified well-formed cotyledons within the central two-thirds of the maternal surface of the placenta. We then cut each cotyledon in half and dissected the histological specimen from the interior of one cut face and the specimen for nucleic acid analysis from the opposite face. The areas sampled consisted mainly of fetal villi and maternal intervillous space. We performed this mirrored sampling on two cotyledons per placenta generating 638 histological and nucleic acid specimens.

Histological specimens were placed in tissue cassettes and fixed with formaldehyde (Merck, purchased in Bamako, the capital city) freshly diluted 1:10 with phosphate buffered saline (PBS) to 3.7% final concentration. PBS was reconstituted using PBS tablets (Sigma-Aldrich) and distilled water. Fixation was for 36 hr. on ice followed by a 30 min. wash in 70% ethanol. The tissue cassettes were shaken off, transferred to plastic bags, and stored in a solar freezer (~ −20 °C) for up to 18 months prior to shipment on dry ice via World Courier to a −80 °C freezer at the University of Michigan. Specimens for nucleic acid analysis were processed as previously described [8].

### 2.2 Slide preparation and visualization

The tissue cassettes were removed from the −80 °C within a year of their arrival and submerged at room temperature in 70% ethanol prior to processing at the University’s Tissue and Molecular Pathology (TMP) core. Tissue blocks were embedded in paraffin and two four-micron thick sections were cut from each sample and stained with hematoxylin and eosin (H&E) or Giemsa. Histology slides were examined under an Olympus BX40 microscope. Polarized light along with location of pigments permitted hemozoin to be distinguished from formaldehyde artifacts [10]. The RNAscope^®^ Multiplex Fluorescent V2 assay (Figure 2) was performed according to the user manual using unstained specimens [11]. RNAscope^®^ slides were imaged on a Leica Stellaris confocal microscope.

We scored histological preservation as 0 for samples without fixation issues, as 1 for samples with damage limited to erythrocytes but still readable, and as 2 if the samples were completely unreadable.

## Results & Discussion

The giemsa-stained specimens (Figure 1) show intact placental morphology, with excellent preservation of the villi, macrophages, neutrophils, and erythrocytes. Figure 1A shows brown hemozoin, the waste product of *P. falciparum*, located in perivillous fibrin of the placenta. Figure 1B shows the gametocyte stage of *P. falciparum* [12], which is the sexual form. Figure 1C shows a severe case of malaria with many *P. falciparum* trophozoites (activated feeding stage). Figure 1D shows a schizont, a mature stage of *P. falciparum*. The asexual stage parasites in A, C, and D indicate active infections. Panel B demonstrates the sexual erythrocytic stage of the parasite that is ingested by mosquitoes during a blood meal [12]. The presence of only hemozoin indicates past infection.

**Fig 1.**
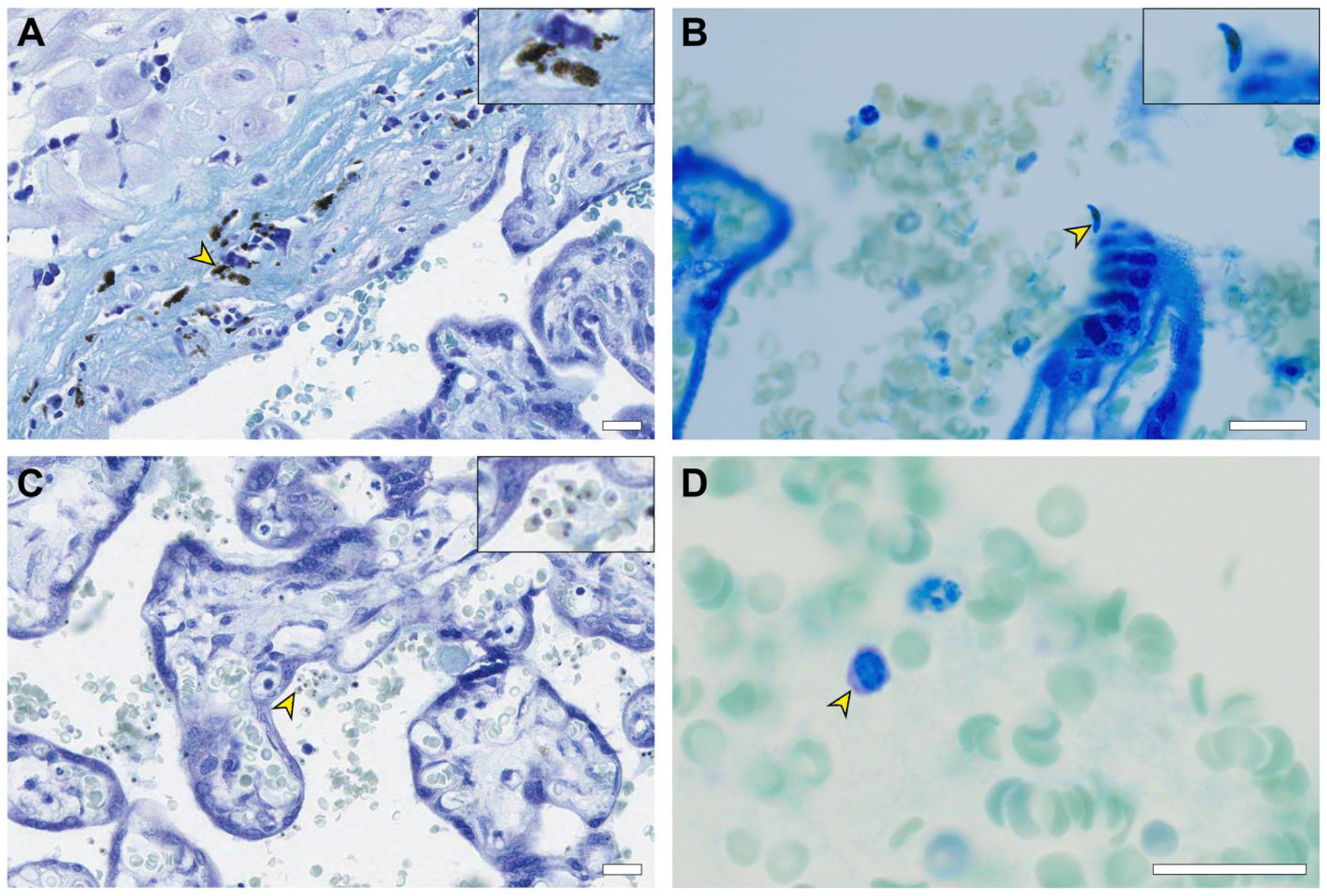
Representative micrographs from specimens of malaria-infected term placentas collected in the field using our novel protocol. Arrows point to features of interest. A shows hemozoin, B shows gametocyte form, C shows trophozoite form, D shows schizont form. Scale bars are 20 μm.

**Figure 2.**
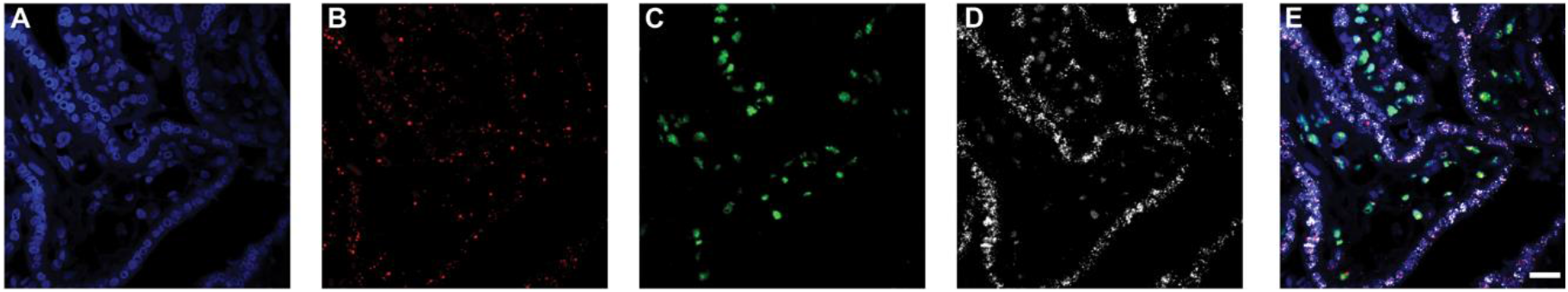
RNAscope^®^ hybridization with probes for three genes. **A.** DAPI **B.** KCNQ1OT1 **C.** MEG3 **D.** ERVW-1 **E.** Merged. Scale bar is 20 μm.

RNAscope^®^ in-situ hybridization was performed to examine whether the preservation of RNA was sufficient to visualize the expression of specific genes. The RNAscope^®^ probes were for two imprinted lncRNAs, KCNQ1OT1 and MEG3, and a protein-coding RNA, ERVW-1, which is a marker for trophoblasts. All three probes generated strong signals in different cells indicative of specific hybridization. Thus, the specimens collected with this protocol were of sufficient quality to provide spatial information on the expression of specific genes.

We collected 638 histology samples from 322 placentas. We excluded 49 due to sample properties unrelated to fixation (e.g. ischemia). Of the remaining 589 samples, 544 (92%) samples were readable (score of 0 or 1). The 45 unreadable specimens could not be assessed for malaria infection due to loss of structure reminiscent of freezing artifacts. We did not attempt to use 30% sucrose as a cryoprotectant due to the inability to assess osmolarity artifacts in the field.

The protocol presented here will improve the collection of placental specimens in low resource settings. Formaldehyde is widely available, but due to its toxicity, appropriate training of personnel is essential. Solar freezers are available in most urban areas and can be installed in rural settings. Maintenance costs consist mainly in the periodic exchange of lead acid batteries. The overall quality of the FFPE specimens was excellent and permitted the evaluation of current and past malaria infection, anemia, placental maturity, and inflammation, as well as visualization of gene expression using RNAscope^®^. The RNA specimens produced high quality data using targeted RNA-seq [13].

## Acknowledgments

This research was supported by the Eunice Kennedy Shriver National Institute of Child Health & Human Development of the National Institutes of Health (R01HD088521 and R21HD077465 to BIS); the John Templeton Foundation (52269 to BIS); the National Science Foundation program in Biological Anthropology (NSF BCS-1354814 to BIS); the Riggs Hoenecke Prize (to PSP); the Michigan Anthropology Undergraduate Grant (to PSP); the Daniel Carl Maier Fund (to PSP); and the EHAP Research Award (to PSP);. The authors thank the study participants who made this research possible.

## Notes

**Declaration of interest:** none

### Competing Interest Statement

The authors have declared no competing interest.

### Summary of Updates

The title was changed to reflect the focus of this study on malaria. The fluorescence micrographs in Fig. 2 have been updated. Authors have been updated as per their request.

## References

[1] R. T. Miller, P. E. Swanson, and M. R. Wick, “Fixation and Epitope Retrieval in Diagnostic Immunohistochemistry: A Concise Review with Practical Considerations,” Appl. Immunohistochem. Mol. Morphol., vol. 8, no. 3, pp. 228–235, Sep. 2000.

[2] J. K. Shetty, H. F. Babu, and K. P. Hosapatna Laxminarayana, “Histomorphological Assessment of Formalin versus Nonformalin Fixatives in Diagnostic Surgical Pathology,” J. Lab. Physicians, vol. 12, no. 04, pp. 271–275, Dec. 2020, doi: 10.1055/s-0040-1722546.

[3] D. G. Vince, A. Tbakhi, A. Gaddipati, R. M. Cothren, J. F. Cornhill, and R. R. Tubbs, “Quantitative Comparison of Immunohistochemical Staining Intensity in Tissues Fixed in Formalin and Histochoice,” Anal. Cell. Pathol., vol. 15, no. 2, pp. 119–129, 1997, doi: 10.1155/1997/607965.

[4] G. J. Burton et al., “Optimising sample collection for placental research,” Placenta, vol. 35, no. 1, pp. 9–22, Jan. 2014, doi: 10.1016/j.placenta.2013.11.005.

[5] J. J. O’Reilly, S. Barak, and A. A. Penn, “A new pipeline for clinico-pathological and molecular placental research utilizing FFPE tissues,” Placenta, vol. 112, pp. 185–188, Sep. 2021, doi: 10.1016/j.placenta.2021.07.301.

[6] P. Pizzolato, “Formalin pigment (acid hematin) and related pigments,” Am. J. Med. Technol., vol. 42, no. 11, pp. 436–440, Nov. 1976.

[7] A. M. Abreu Velez, M. S. Howard, M. Restrepo-Isaza, and B. Smoller, “Formalin deposition as artifact in biopsies from patients affected by a new variant of endemic pemphigus foliaceus in El Bagre, Colombia, South America,” J. Cutan. Pathol., vol. 37, no. 8, pp. 835–842, 2010, doi: 10.1111/j.1600-0560.2009.01492.x.

[8] C. Vincenz, J. L. Lovett, W. Wu, K. Shedden, and B. I. Strassmann, “Loss of Imprinting in Human Placentas Is Widespread, Coordinated, and Predicts Birth Phenotypes,” Mol. Biol. Evol., vol. 37, no. 2, pp. 429–441, Feb. 2020, doi: 10.1093/molbev/msz226.

[9] B. I. Strassmann, “Cooperation and competition in a cliff-dwelling people,” Proc. Natl. Acad. Sci., vol. 108, no. Supplement_2, pp. 10894–10901, Jun. 2011, doi: 10.1073/pnas.1100306108.

[10] C. Romagosa et al., “Polarisation microscopy increases the sensitivity of hemozoin and Plasmodium detection in the histological assessment of placental malaria,” Acta Trop., vol. 90, no. 3, pp. 277–284, May 2004, doi: 10.1016/j.actatropica.2004.02.003.

[11] F. Wang et al., “RNAscope: A Novel in Situ RNA Analysis Platform for Formalin-Fixed, Paraffin-Embedded Tissues,” J. Mol. Diagn., vol. 14, no. 1, pp. 22–29, Jan. 2012, doi: 10.1016/j.jmoldx.2011.08.002.

[12] R. Tuteja, “Malaria - an overview,” FEBS J., vol. 274, no. 18, pp. 4670–4679, 2007, doi: 10.1111/j.1742-4658.2007.05997.x.

[13] W. Wu, J. L. Lovett, K. Shedden, B. I. Strassmann, and C. Vincenz, “Targeted RNA-seq improves efficiency, resolution, and accuracy of allele specific expression for human term placentas,” G3 GenesGenomesGenetics, vol. 11, no. 8, p. jkab176, Aug. 2021, doi: 10.1093/g3journal/jkab176.

